# Host origin of microbiota drives functional recovery and *Clostridioides difficile* clearance in mice

**DOI:** 10.1101/2024.11.19.624317

**Authors:** Sophie A. Millard, Kimberly C. Vendrov, Vincent B. Young, Anna M. Seekatz

**Author notes:** Corresponding author: Contact information: Anna M. Seekatz, PhD, ^a^Department of Biological Sciences, Clemson University, Clemson, SC 29634, USA. **Patient consent:** Informed consent was obtained from individuals prior to the time of sampling. IRB approved on 02/04/2015.

## Abstract

Colonization resistance provided by the gut microbiota is essential for resisting both initial *Clostridioides difficile* infection (CDI) and potential recurrent infection (rCDI). Although fecal microbiota transplantation (FMT) has been successful in treating rCDI by restoring microbial composition and function, mechanisms underlying efficacy of standardized stool-derived products remain poorly understood. Using a combination of 16S rRNA gene-based and metagenomic sequencing alongside metabolomics, we investigated microbiome recovery following FMT from human and murine donor sources in a mouse model of rCDI. We found that a human-derived microbiota was less effective in clearing *C. difficile* compared to a mouse-derived microbiota, despite successful microbial engraftment and recovery of bacterial functional potential. Metabolomic analysis revealed deficits in secondary metabolites, suggesting a functional remodeling between human microbes in their new host environment. Collectively, our data revealed additional environmental, ecological, or host factors involved in FMT-based recovery from rCDI.

**Importance:** *Clostridioides difficile* is a significant healthcare-associated pathogen, with recurrent infections presenting a major treatment challenge due to further disruption of the microbiota after antibiotic administration. Despite the success of fecal microbiota transplantation (FMT) for the treatment of recurrent infection, the mechanisms mediating its efficacy remain largely underexplored. This study reveals that effectiveness of FMT may be compromised by a mismatch between donor microbes and the recipient environment, leading to deficits in key microbial metabolites. These findings highlight additional factors to consider when assessing the efficacy of microbial-based therapeutics for CDI and other conditions.

## Introduction

Colonization resistance against the healthcare-associated pathogen, *Clostridioides difficile,* is predominantly mediated by the indigenous microbes in our gastrointestinal tract, termed the gut microbiota (1). A diverse gut microbiota typically helps prevent initial colonization and disease if an individual encounters *C. difficile* spores from the environment (2, 3). However, disruptions to this ecosystem, such as following antibiotic treatment (4–7), can create conditions conducive to *C. difficile* spore germination, outgrowth, and toxin production, leading to *C. difficile* infection (CDI) (8, 9).

Given the crucial role of the microbiota in preventing CDI, there is significant interest in developing alternatives to standard antibiotic therapies directed against *C. difficile*, which can further disrupt the microbiota and heighten the risk of recurrent CDI (rCDI) (10, 11). Antibiotics that specifically target *C. difficile* while preserving the gut microbiota show promise in reducing rCDI rates (12, 13). Advances in treating rCDI have been demonstrated by the success of microbial-based therapeutics, notably fecal microbiota transplantation (FMT), which involves transferring stool from a healthy donor to a diseased individual to restore the recipient’s microbiota (14). In 2023, the FDA approved two standardized stool-derived products for rCDI treatment (15, 16). While these products have mitigated some safety concerns associated with using minimally processed stool, they remain non-specific and still carry safety and efficacy concerns (17–19). Despite the extensive research demonstrating potential mechanisms by which gut microbes inhibit *C. difficile* (20–24), single probiotic or targeted microbial formulations do not always achieve the success rates of FMT or stool-derived products (25, 26). Consequently, a significant gap in knowledge remains regarding the specific factors and interactions that determine long-term stability and success of the transplanted microbiota.

Both engraftment of specific microbial species and restoration of overall microbial diversity is correlated with successful FMT outcomes (27–30). Mechanistically, these species are hypothesized to reconstitute bacterial functions that inhibit *C. difficile*. One key microbial function associated with FMT success is the transformation of primary bile acids in the gut, which is exclusively performed by microbes and is known to inhibit vegetative growth of *C. difficile* (20, 31, 32). Additionally, the levels of the bacterial fermentation products, such as the short chain fatty acid (SCFA) butyrate, have been positively correlated with successful FMT outcomes in both mouse studies and patients with rCDI (33, 34). In addition to microbial-derived metabolites, recent studies have also emphasized the importance of bacterial nutrient exclusion in limiting *C. difficile* colonization (35–38). *C. difficile* is auxotrophic for several amino acids, such as proline (39, 40), and can ferment amino acids, which are more abundant in a gut with reduced bacterial diversity (5). Transplanting a microbiota capable of competing with *C. difficile* for these nutrients likely plays a crucial role in successful clearance. The host environment may also play a role in FMT success in addition to microbial functional restoration. For instance, the host immune response has been observed to influence the ability of FMT to clear *C. difficile* in mice (41). Increased inflammation can also change *C. difficile* metabolism and virulence (42), as well as microbiota interactions (22, 43), that could influence FMT outcome. Understanding these additional interactions when transplanting microbial communities between hosts is thus a critical consideration for therapeutics relying on microbiota manipulation, particularly for conditions beyond *C. difficile* where success rates are not as high (44, 45).

This study aimed to identify functions important for clearing *C. difficile* in the gut. However, our results also highlighted additional factors to consider when translating microbial community structure into functional outcomes, particularly in the context of host-microbe adaptation in the gut. Using a mouse model of rCDI, we observed that compared to a mouse-derived microbial community, a diverse human-derived microbial community was unable to clear *C. difficile*. Independent of their ability to clear *C. difficile*, both fecal products demonstrated engraftment of microbes typically associated with successful FMT outcome. Metagenomic sequencing also demonstrated recovery of bacterial genetic functions important for *C. difficile* clearance, suggesting recovery of functional potential, independent of *C. difficile* clearance. In contrast, both untargeted and targeted metabolomics demonstrated deficits in many secondary metabolites, in line with previous metabolomic comparisons before and after FMT. Collectively, these results suggest human microbes, when transplanted into a mouse with an altered murine microbiome, are unable to realize their functional potential in this new host environment. These results underscore the need to consider not only the functional potential of a microbial-derived therapeutic, but how to ensure that these functions manifest in a treated patient.

## Methods

### Ethics Statement

All animal protocols were approved by the University of Michigan Institutional Animal Care and Use Committee (protocol # PRO00008114), which adhere to the Public Health Service Policy on Humane Care and Use of Laboratory Animals guidelines. Informed consent was obtained from individuals prior to fecal donation using protocols approved by the University of Michigan Institutional Review Board (#HUM00130242, #HUM00098164).

### Mouse model of recurrent CDI

All experiments used 5 – 8 week-old C57BL/6 male and female mice from an established breeding colony at the University of Michigan, originally sourced from Jackson Laboratories (Bar Harbor, ME). Animal housing was conducted in specific-pathogen-free and biohazard (autoclave-in / autoclave-out) conditions, received autoclaved food, water, and bedding in a 12-hour light/dark cycle. A laminar flow hood with personal protective equipment and the use of the sporicidal disinfectant Perisept (Triple S, Navigator #62, Los Angeles, CA) was used for all cage changes, infections, and sample collections. Mice were housed in groups of 3 – 5 animals per cage, ensuring multiple cages per group. Results represent eight sets of experiments, with cage assignments and experimental groups detailed in Table S1.

A previously described CDI recurrence model was used for all experiments (46). Mice (n = 129) were administered 0.5 mg / ml of cefoperazone (MP Biochemicals, #199695) in sterile drinking water (Gibco, #15230) for 5 days. At day 0, mice received 10^3^ spores of *C. difficile* strain 630 (ATCC BAA-1382) in 20 μl sterile PBS (Gibco, #10010) via oral gavage, generated as previously described (47). Mice were given 0.4 mg/ml vancomycin (Sigma, #V2002; #V8138) in drinking water days 4 – 9. At day 11, mice were left untreated (‘noFMT’; n = 24) or administered a 100 μl fecal preparation via oral gavage from one of the following sources: feces from healthy, untreated, age-matched animals from the same breeding colony (‘mFMT’; n = 33), feces from healthy, untreated animals from other breeding colonies (‘mFMT-other’; n = 23), or feces from one of six human donors (‘hFMT’; n = 53). Detailed preparation protocols and donor sources are further described in Supplemental Methods. Mice were monitored daily for clinical signs of CDI and weight loss, with indicated fecal samples collected directly from mice throughout the experiment and cecal content collected at euthanization at early (day 21) or late (day 42) timepoints.

Fecal and cecal samples were enumerated in an anaerobic chamber (Coy Laboratory Products, Grass Lake, MI) for *C. difficile* via colony-forming units (CFUs) (46). Briefly, samples were homogenized in pre-reduced PBS in a 1:10 ratio based on sample weight, and serially diluted to 10^-6^. Multiple 100 μl dilutions were plated onto taurocholate cycloserine cefoxitin fructose agar (TCCFA) plates (48) for overnight incubation prior to CFU enumeration.

### DNA extraction, library preparations, and sequencing

DNA was extracted with the MO Bio PowerFecal kit (now PowerFecalDNA, Qiagen, Hilden Germany), adapted to the epMotion 5075 TMX (Eppendorf, Hamburg, Germany). The UMICH Microbiome Core conducted all DNA library preparation and 16S rRNA gene-based sequencing (46), as developed by Kozich et al (48). Briefly, the V4 region of the 16S rRNA gene was PCR-amplified using barcoded dual-index primers. Upon confirmation of a correctly sized PCR product using gel electrophoresis (Invitrogen, #G401002), PCR products were normalized using the SequelPrep plate kit (Life Technologies, #A10510-01) and pooled per 96-well plate. Each pool was quantified using qPCR (KapaBiosystems, #KK4854) and sized using the Agilent Bioanalyzer high-sensitivity DNA kit (Agilent, #5067-4642). The Illumina MiSeq platform with the MiSeq Reagent 222 kit v2 (#MS-102-2003) was used to sequence amplicons with a 10% PhiX spike according to manufacturer’s protocol using a final concentration of 4 pM.

Metagenomic sequencing was conducted by the University of Minnesota Genomics Center (UMGC) and the Clemson University Genomics and Bioinformatics Facility (CUGBF) (Table S1). All libraries were prepared with Nextera XT DNA Library Prep kit (part #15032355), quantified using the Kapa qPCR, and sized via the Agilent Bioanalyzer before pooling to an equimolar concentration and sequencing, using paired-end 2×150 settings on either the Illumina NovaSeq 6000 or NextSeq 550 platform.

### 16S rRNA gene-based analyses

Sequences were processed using mothur v1.37.6 (49), with specific quality parameters and commands indicated in the associated data repository (‘Code Availability’). Briefly, the SILVA rRNA database project (v128) (50) was used to align reads to the V4 region of the 16S rRNA gene, using UCHIME to remove chimeric sequences (51). Sequences were taxonomically classified using the mothur-adapted version of the RDP database (v16) (52) using the Wang method (80% minimum bootstrap) (53). Operational taxonomic units (OTUs) were clustered to 97% similarity using the OptiClust algorithm in mothur (54) and used for Shannon diversity index and pairwise Bray-Curtis dissimilarity index values. A combination of base R commands and packages were used for data visualization and statistical analyses. Nonmetric multi-dimensional scaling (NMDS) and PERMANOVA were implemented using vegan (55). The Kruskal-Wallis test was used for statistical significance across multiple groups, with a post-hoc Dunn’s test when applicable. Multivariable Association with Linear Models (MaAsLin2) was used to calculate significantly abundant OTUs between cleared vs colonized mice (56).

### Metabolomics

Detailed processing and quantification steps are described in Supplemental Methods. Briefly, untargeted metabolomics from cecal samples were conducted by Metabolon (Durham, NC) using Ultrahigh Performance Liquid Chromatography-Tandem Mass Spectroscopy (UPLC-MS/MS) (Table S3). Targeted cecal measurements of bile acids (LC-MS) and SCFAs (GC-MS) were conducted by the University of Michigan Metabolomics Core. Fecal SCFA measurements (HPLC) were conducted at the University of Michigan Microbiome Core, as previously described (57).

### Metagenomic taxonomic and functional profiling

Host reads and adapters were removed from raw reads using KneadData (v0.12.0). Quality-filtered metagenomes were taxonomically profiled using MetaPhlAn v4.1 using the vOct22 CHOCOPhlanSGB with default parameters (58).

Functional gene annotation and quantification of filtered sequence data was conducted using HUMAnN v3.6 (59) against the Uniref90 functional gene database (60) with default settings. Pathway annotation was also conducted using HUMAnN3 against the MetaCyc v24.0 database (61). Generated reads per kilobase per million (RPKM) counts of annotations were normalized to copies per million (CPM) to account for sequencing depth variability. For global assessment of genes, Uniref90 IDs were regrouped as KEGG orthologs (KO) (62) using the *humann_regroup_tables* command. A combination of base R commands and packages were used for data visualization and statistical analyses. Inverse-simpson diversity, pairwise Bray-Curtis dissimilarity index values, nonmetric multi-dimensional scaling (NMDS), and PERMANOVA were implemented using vegan (55). Significant differences in taxonomic or gene abundance was determined using MaAsLin2 package (56). The Kruskal-Wallis test was used for statistical significance across multiple groups, with a post-hoc Wilcoxon rank-sum or Dunn’s test when applicable.

### Data and code availability

Raw sequence data have been deposited in the Sequence Read Archive (Project PRJNA1168499). All code involved in generating analyses for this study, including processed raw data, is available at https://github.com/SeekatzLab/mouseCDI-SPF-hFMT.

## Results

### Human-derived microbiota engraft in mice but fail to clear C. difficile in a mouse model of recurrent infection

We previously developed a mouse model of rCDI, demonstrating that employing mouse-FMT derived microbiota from healthy, untreated mice (mFMT) rapidly cleared *C. difficile* (46, 63). In this model, mice are rendered susceptible to *C. difficile* with cefoperazone prior to spore inoculation (Figure 1A). At maximal disease severity (day 4), mice are treated with vancomycin, which results in *C. difficile* suppression. However, upon vancomycin cessation, *C. difficile* will re-colonize. In the current study, we aimed to identify human-specific microbiota with the capacity to clear *C. difficile* and used fecal material from different healthy human donors, including a subset of fecal sources previously used to successfully treat human patients (hFMT; n = 6 unique fecal samples) (Table S1) (34). As previously observed, mice lost weight during initial and recurrent infection (Figure 1B). Without treatment (noFMT), mice remained colonized (Figure 1C). Successful clearance of *C. difficile* was once again observed by healthy mouse feces (mFMT), as well as additional mouse-derived fecal material from other genotypes colonies or a spore-preparation of mouse feces (mFMT-other; Table S1 and Methods). In contrast to mFMT, none of the hFMT sources used in our study demonstrated capacity to clear *C. difficile* (Figure 1C, D; Table S1) (29).

**Figure 1.**
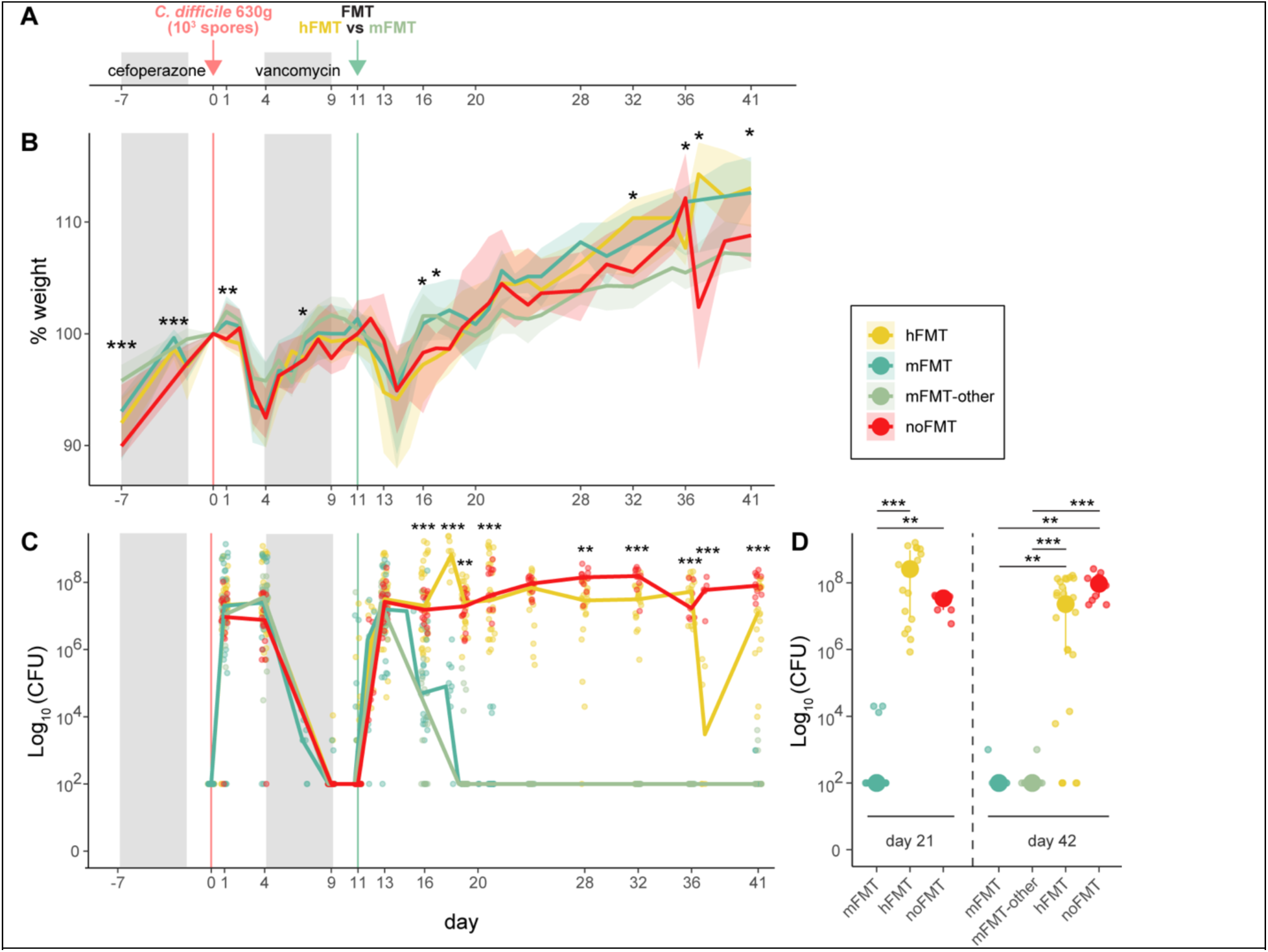
Human-derived fecal material fails to clear *C. difficile* in mice with recurrent *C. difficile* infection (CDI). **A)** Mouse model of recurrent CDI. **B)** Percent weight loss (compared to day 0, day of *C. difficile* infection) over time in mice treated with feces from healthy humans (hFMT; n = 53), colony-matched mice (mFMT; n = 33), mice representative of different genetic or colony backgrounds (mFMT-other; n = 23), or untreated mice (noFMT; n = 24). *C. difficile* colonization, as assessed using the log_10_-normalized colony-forming units (CFU), identified in **C)** fecal samples over time or **D)** cecal samples at day 21 or day 42 post-infection in mice treated with hFMT, mFMT, hFMT, or untreated (noFMT) mice. Statistical significance determined using Kruskal-Wallis test, with a post-hoc Dunn test for **(D)** (\**p* < 0.01, \*\**p* < 0.001, \*\*\**p* < 0.0001).

We conducted 16S rRNA gene-based sequencing to identify whether failure to clear *C. difficile* was due to limited engraftment of human-derived microbiota in recipient mice. As assessed by non-metric multidimensional scaling (NMDS) of the Bray-Curtis dissimilarity calculated from operational taxonomic units (OTUs), the overall microbiota structure was shifted following both antibiotic treatments and *C. difficile* inoculation (PERMANOVA, *p* < 0.001; Figure 2A). Although fecal specimens from hFMT recipients were less similar to their input communities than either mFMT or mFMT-other (Dunn’s test, *p* < 0.0001; Figure 2B, left panel), variability across recipients in each treatment group was relatively similar, with the exception of mice given mFMT-other, which were more similar to each other (Dunn’s test, *p* < 0.0001; Figure 2B, middle panel). Inter-group dissimilarity was high across all recipient comparisons, with mice receiving mFMT versus noFMT displaying the highest dissimilarity (Figure 2B, right panel).

**Figure 2.**
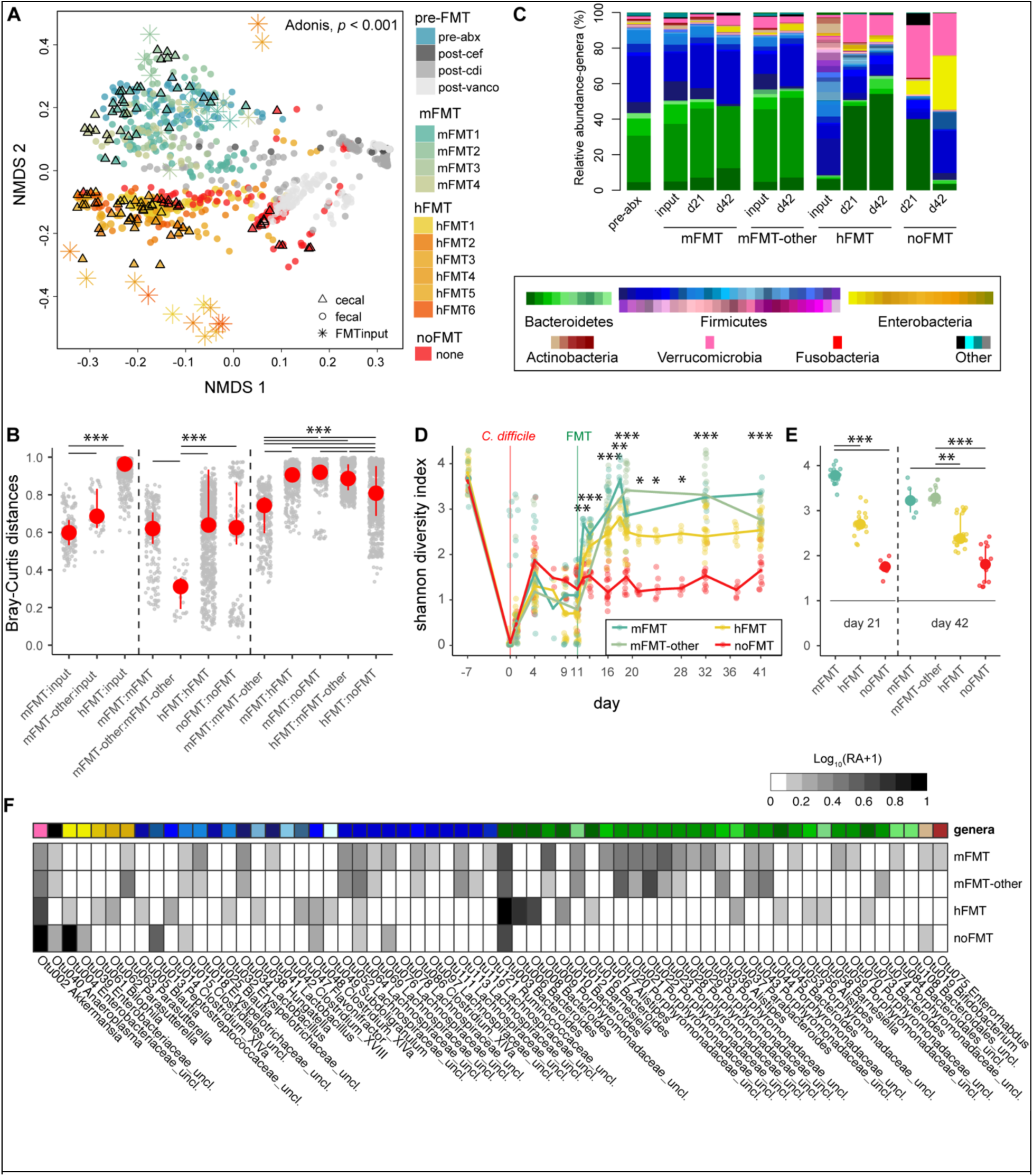
Human-derived fecal bacteria engraft mice with CDI. **A)** Non-metric multidimensional (NMDS) scaling of the Bray-Curtis dissimilarity index calculated from operational taxonomic units (OTUs) throughout CDI timeline. Fecal samples throughout the experiment or endpoint cecal samples were collected throughout experiment (pre-FMT) without treatment (noFMT) and following treatment with various mouse (mFMT1 – 4) or human (hFMT1 – 6) feces, with sample designation (including FMT inputs) as indicated in legend. PERMANOVA, *p* < 0.001 calculated across groups (mFMT, hFMT, and noFMT) for both cecal and fecal timepoints. **B)** Pairwise Bray-Curtis dissimilarity calculated between treatment groups (mFMT, mFMT-other, hFMT, and noFMT) and their respective inputs or between and within treatment groups. **C)** Average relative abundance of top 98% genera observed in cecal samples prior to any treatment (pre-abx) or at indicated timepoints without treatment (noFMT) or following mFMT, mFMT-other, or hFMT. Shannon diversity index calculated from **C)** fecal samples over time or **D)** cecal samples at day 21 or day 42 post-infection in mice treated with hFMT, mFMT, hFMT, or untreated (noFMT) mice. **F)** Differentially abundant OTUs between mice that did (mFMT, mFMT-other) versus did not clear (hFMT, noFMT) *C. difficile* (MaAsLin2, linear model with BH correction, *q* ≤ 0.001) with the log_10_-normalized relative abundance (RA+1) per OTU per treatment group. Statistical significance for **(B, D, E)** determined using Kruskal-Wallis test, with a post-hoc Dunn test for **(B, E)** (\**p* < 0.01, \*\**p* < 0.001, \*\*\**p* < 0.0001).

Composition of the microbiota across treatments demonstrated increased members of Firmicutes (Bacillota) and Bacteroidetes (Bacteroidota) in both hFMT and mFMT-treated mice compared to the noFMT group, which was primarily dominated by Enterobacteria and Verrucomicrobia (Verrucomicrobiota) (Figure 2C). Specifically, many genera frequently associated with resistance to or recovery from *C. difficile* in mouse and human studies (30, 34, 64) were increased following both hFMT and mFMT, including *Bacteroides* and unclassified Lachnospiraceae species (Figure 2D, Figure S1). Within Bacteroidetes, *Bacteroides* dominated mice given hFMT whereas mouse-specific unclassified Porphyromonadaceae and *Alistipes* were more abundant in mice given mFMT. Within Firmicutes, *Lactobacillus*, unclassified Clostridiales, and unclassified Peptostreptococcaceae (inclusive of *C. difficile*) were more predominant in noFMT mice, whereas a variety of Firmicutes genera, including *Blautia*, *Faecalibacterium*, and unclassified Lachnospiraceae species, were observed in mFMT and hFMT groups. Mice given hFMT also demonstrated increased overall diversity compared to noFMT mice, although not as high as mice given mFMT (Dunn’s test, *p* < 0.0001; Figure 2D, E). Using MaAsLin2, we identified several differentially abundant OTUs between mice that did (mFMT, mFMT-other) or did not (hFMT, noFMT) clear. However, comparison of the abundance of these OTUs across all four groups demonstrated that many OTUs in mice given mFMT were still taxonomically represented within mice given hFMT, suggesting the presence of different but taxonomically similar species in both FMT groups. Collectively, these results support the engraftment of human-adapted microbiota in mice, including genera typically associated with successful treatment of rCDI via microbiota replacement.

### FMT input source dictates species-level composition post-transplantation independent of C. difficile clearance

We conducted metagenomic sequencing to further resolve taxonomic differences between the treated mice. Similar to the 16S rRNA gene-based taxonomic profiles, the ceca of mice treated with either FMT exhibited distinct strain-level composition from untreated mice, as demonstrated by NMDS of the Bray-Curtis dissimilarity based on species abundance using MetaPhlan4 (PERMANOVA, *p* < 0.001; Figure 3A). Minor clustering by individual hFMT donors was also observed, although not as pronounced as among the three treatment groups. Bray-Curtis distance was highest between hFMT- and mFMT-treated mice, although the other comparisons (mFMT:noFMT, hFMT:noFMT) were also significantly different (Dunn’s test, *p* < 0.0001; Figure 3B). Compared to mice that received mFMT, untreated mice exhibited significantly decreased species level diversity (Wilcoxon rank-sum test, p < 0.05; Figure 3C). Decreased diversity, albeit higher than untreated mice, was also observed in mice treated with hFMT (Wilcoxon rank-sum test, *p* < 0.05; Figure 3C). Mice treated with either FMT were dominated by diverse Firmicutes and Bacteroidetes species compared to untreated mice, which exhibited an expansion of Verrucomicrobia (Figure 3D). Individual variation between microbiota of the mice was observed, although overall trends in composition clustered by FMT source (Supplementary S2).

**Figure 3.**
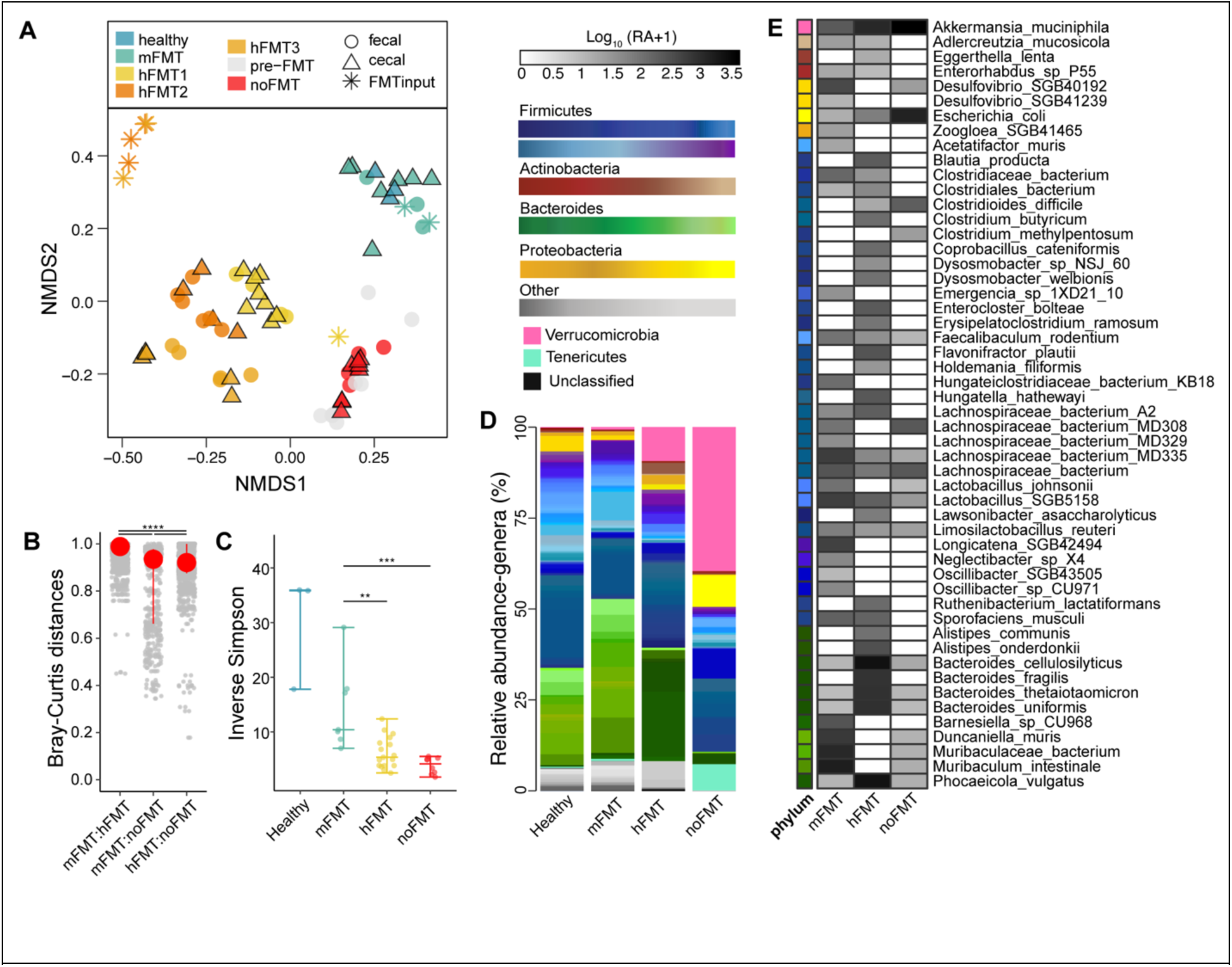
FMT source determines species composition post-transplantation, regardless of *C. difficile* clearance. **A)** NMDS of Bray-Curtis dissimilarity (stress= 0.1390) and **B)** Bray-Curtis distances based on bacterial species identified in fecal and cecal samples from mice. Asterisks denote significance (**** *p* < 0.0001) determined using Kruskal-Wallis test, with a post-hoc Dunn test. **C)** Inverse Simpson Diversity of bacterial species identified in cecal samples. Asterisks denote significance (** *p* < 0.05) determined using Kruskal-Wallis test, with a post-hoc pairwise Wilcoxon rank-sum test. **D)** Average relative abundance of bacterial genera identified by MetaPhlAn4 from cecal samples of mice. **E)** Mean log_10_-transformed (CPM + 1) of named bacterial genera significantly different in abundance across treatment groups compared to mice treated with mFMT (based on MaAsLin2; linear model with BH correction, *q* ≤ 0.01).

MaAsLin2 was used to identify differentially abundant bacterial taxa between mice that did (“cleared”; mFMT) or did not clear *C. difficile* (“colonized”; hFMT or noFMT). Many of the bacterial species with the largest variation were unnamed members of the Firmicutes phylum (linear model with BH correction; q ≤0.001; Supplementary S3). Several species identified, including *Acetatifactor muris*, *Faecalibaculum rodentium*, and *Duncaniella muris,* are commonly found in mouse microbiota (65, 66) and are not typically associated with *C. difficile* infection or clearance (Figure S3). We also used MaAsLin2 to identify differentially abundant species across all three groups, independent of *C. difficile* clearance (Supplementary S4). This multi-group comparison demonstrated taxa unique to each group, although the majority were unnamed or unclassified (linear model with BH correction, *q* ≤ 0.01; Supplementary S4). Among the 165 species identified as differentially abundant, only 52 were named at the species level (linear model with BH correction, *q* ≤ 0.01; Figure 3E). Species such as *Clostridium butyricum*, *Flavonifractor plautii*, and *Eggerthella lenta* were unique to hFMT-treated mice, whereas only *Clostridium methylpentosum* was unique to untreated mice (Figure 3E). Some species that increased in hFMT-treated mice compared to mFMT-treated mice were those associated with a typical human microbiota, including *Bacteroides fragilis*, *Bacteroides thetaiotaomicron*, and *Phoecaciola vulgatus* (Figure 3E). Collectively, our results suggest similar findings from initial 16S rRNA gene-based sequencing, in that human-derived taxa colonize mice post-FMT despite their inability to clear *C. difficile*.

### Metagenomic functions vary depending on source of FMT, independent of C. difficile clearance

To investigate the functional potential of the engrafted microbiota, we used Humann2 to identify KEGG Ortholog (KO) counts from the metagenomes. Based on Bray-Curtis dissimilarity of normalized KO abundances, the gene-encoded functional profile also demonstrated distinct clustering by treatment group, with some mice that received hFMT exhibiting overlapping KO profiles with mice that received mFMT (PERMANOVA, *p* < 0.001; Figure 4A). Confirming what was seen on with NMDS, Bray-Curtis distances among all comparisons showed that hFMT samples were significantly more dissimilar to noFMT samples than mFMT samples (Dunn’s test, *p < 0.05*; Figure 4B). We observed 143 over-represented KOs and 170 underrepresented KOs in mice that did not clear *C. difficile* compared to those that did (linear model with BH correction, *q* ≤ 0.001; Figure 4C). The majority of these (n=160) were related to metabolism (Figure S5), with the remainder (n=153) classified under genetic information processing, cellular processing, or environmental processing (Figure S6). For metabolism-related KOs, several were more abundant in the microbiota of mice that did not clear *C. difficile*. These included genes potentially involved in detoxifying inflammatory products derived from the host such as glutaminase, catalase, and those involved in ethanolamine utilization (Figure S5) Additionally, several translation-associated genes were more abundant in the microbiota of these mice (Figure S6). Conversely, KOs more abundant among the microbiota of mice that cleared *C. difficile* include those involved in urea metabolism and metal nutrient uptake (Figure S5, S6).

**Figure 4.**
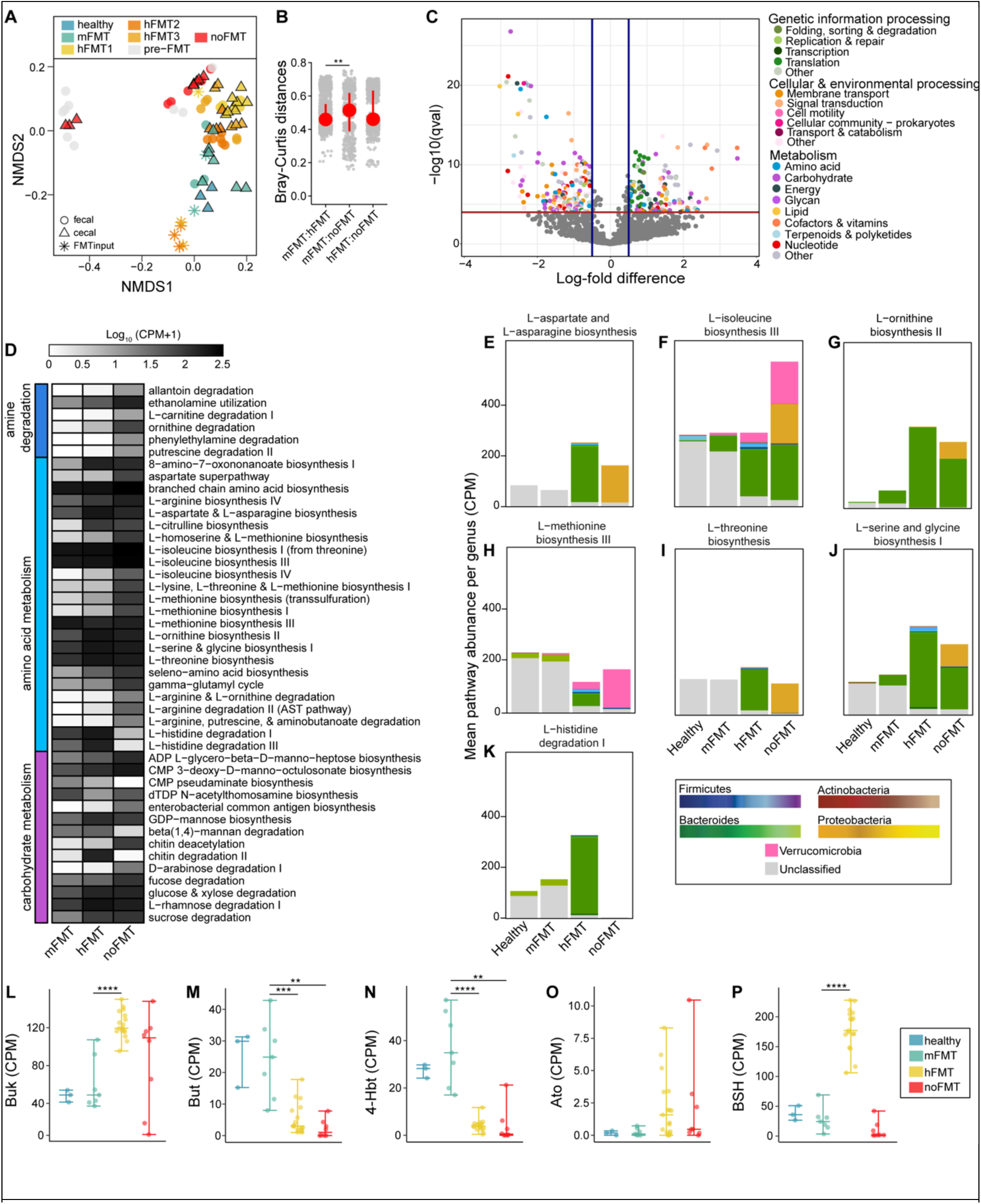
Genes, pathways, and contributing taxa vary with FMT source, independent of *C. difficile* clearance. **A)** NMDS of Bray-Curtis dissimilarity (stress= 0.1109) and **B)** Bray-Curtis distances based on KEGG Orthologs (KOs) identified in fecal and cecal samples from mice. Asterisks denote significance (** *p* < 0.05) determined using Kruskal-Wallis test, with a post-hoc Dunn test **C)** KOs identified to be significantly different in abundance between mice that did or did not clear *C. difficile* (based on MaAsLin2; linear model with BH correction, *q* ≤ 0.001). **D)** Mean log_10_-transformed (CPM + 1) of amine, amino acid, and carbohydrate metabolism MetaCyc Pathways significantly different across treatment groups compared to mice treated with mFMT (based on MaAsLin2; linear model with BH correction, *q* ≤ 0.001). **E-K)** CPM of genes belonging to the labeled amino acid metabolism MetaCyc Pathway, colored by the genus that encodes it. **L-P)** CPM of Uniref90 genes associated with bile salt hydrolase (*Bsh*) or terminal genes involved in butyrate production (*buk, but, 4-Hbt,* and *Ato*). Asterisks denote significance (** *p* < 0.01, *** *p* < 0.001, **** *p* < 0.0001) determined using Kruskal-Wallis test, with a post-hoc pairwise Wilcoxon rank-sum test.

Due to limitations with KEGG annotation for certain microbial metabolic functions, we used the Uniref90 database to investigate whether the abundance of genes previously associated with *C. difficile* differed across groups. We hypothesized that *bsh*, the gene responsible for deconjugating primary bile acids, would be over-represented in mice that cleared *C. difficile*, given the importance of bile acids in *C. difficile* pathogenesis (21, 23, 24). Strikingly, hFMT-treated mice had the highest abundance of this gene (Wilcoxon rank-sum test, *p* < 0.0001; Figure 4P). We did not detect a complete *bai* operon, required for 7α-dehydroxylation of unconjugated bile acids (67) of in any treatment group (Figure S7). We also looked at the abundance of genes involved in the final step of butyrate production (68), which has been correlated with successful FMT treatment (34) and clearance of *C. difficile* (33) (Figure 4C). Of the main butyrate-producing genes, butyryl-CoA:acetate CoA transferase (*but*) and butyrate kinase (*buk*), *but* was increased in mice given mFMT (Wilcoxon rank-sum test, *p* < 0.001; Figure 4M), but *buk* was increased in mice given hFMT and untreated mice (Wilcoxon rank-sum test, *p* < 0.0001; Figure 4L). A similar pattern was observed for two additional genes that encode terminal enzymes involved in producing butyrate from alternative substrates, succinate (*4hbt*) and lysine (*ato*), where *4hbt* was more abundant in mice that cleared (mFMT) (Wilcoxon rank-sum test, *p* < 0.00001; Figure 4N). Although not significant, *ato* was more abundant in mice that did not clear *C. difficile* (Figure 4O).

Given the gene-level functional differences across the microbiomes of mice from different treatment groups, we expanded our analysis to explore these variations at a pathway level. Using MaAsLin2, we identified MetaCyc pathways with significant differences in abundance between mice that received hFMT or no FMT and those receiving mFMT (linear model with BH correction, *q* ≤ 0.001; Figure 4D, S8). We identified numerous amine, amino acid, and carbohydrate pathways present across the microbiota of mice in all treatment groups, with distinct variation in pathway abundance depending on the type of FMT administered (Figure 4D). In mice that received no treatment, several pathways involved in amino acid metabolism, including some for which *C. difficile* is auxotrophic for, were present at higher abundance (Figure 4D).

Recognizing the importance of amino acid utilization in *C. difficile* clearance, we next examined whether the taxa contributing these amino acid pathways would be different (Figure 4E-K). In the microbiota of mice that received no FMT, these pathways were dominated by Proteobacteria (Pseudomonadota), whereas the pathways of hFMT-treated mice were predominately attributed to Bacteroides and Firmicutes members (Figure 4E-K). Interestingly, the microbiota contributing these functions in both healthy and mFMT-treated mice are largely unclassified (Figure 4E-K). Altogether, our results reveal distinct functional differences in the microbiota of mice based on FMT treatment, with key genes and pathway differences related to detoxification, amino acid metabolism, and metabolite production.

### Mice treated with human-derived microbiota have limited restoration of microbial metabolites typically associated with C. difficile clearance

While recovery of the microbiota community structure is correlated with CDI recovery, it is the functional recovery provided by this community that ultimately contributes to resistance or clearance. To compare how mFMT and hFMT impacted the realized functions in the gut after FMT, we conducted untargeted metabolomics of a subset of cecal samples. This included samples from mice given mFMT or hFMT (from three different donors) compared to no treatment. Comprehensive differences among the three groups were observed (PERMANOVA, *p* < 0.001), as assessed by NMDS of the Bray-Curtis dissimilarity based on the median-scaled and minimum-imputed abundance of metabolites (Figure 5A). Based on pairwise Bray-Curtis distance calculations, mice receiving mFMT were significantly more similar to healthy mice than any other group (Dunn’s test, *p* < 0.001; Figure 5B, right panel). Intra-group dissimilarity was lower than inter-group dissimilarity, with the highest dissimilarity observed between mice receiving mFMT compared to mice receiving no treatment (Figure 5B, middle and right panels). Random Forest analysis of the most important features between cleared (healthy or mFMT-treated mice) and colonized mice (hFMT or noFMT) identified increased lipid-classified compounds in cleared animals, including the bile acids isohyodeoxycholate and taurohyodeoxycholate, and the SCFAs butyrate and valerate (Figure 5C). In contrast, many amino acids and carbohydrates were decreased in cleared animals, including polyamines such as N-acetyl-cadaverine and N-acetyleputrescine and neuropeptides such as N-Acetylaspartylglutamic acid and gamma-aminobutyric acid.

**Figure 5.**
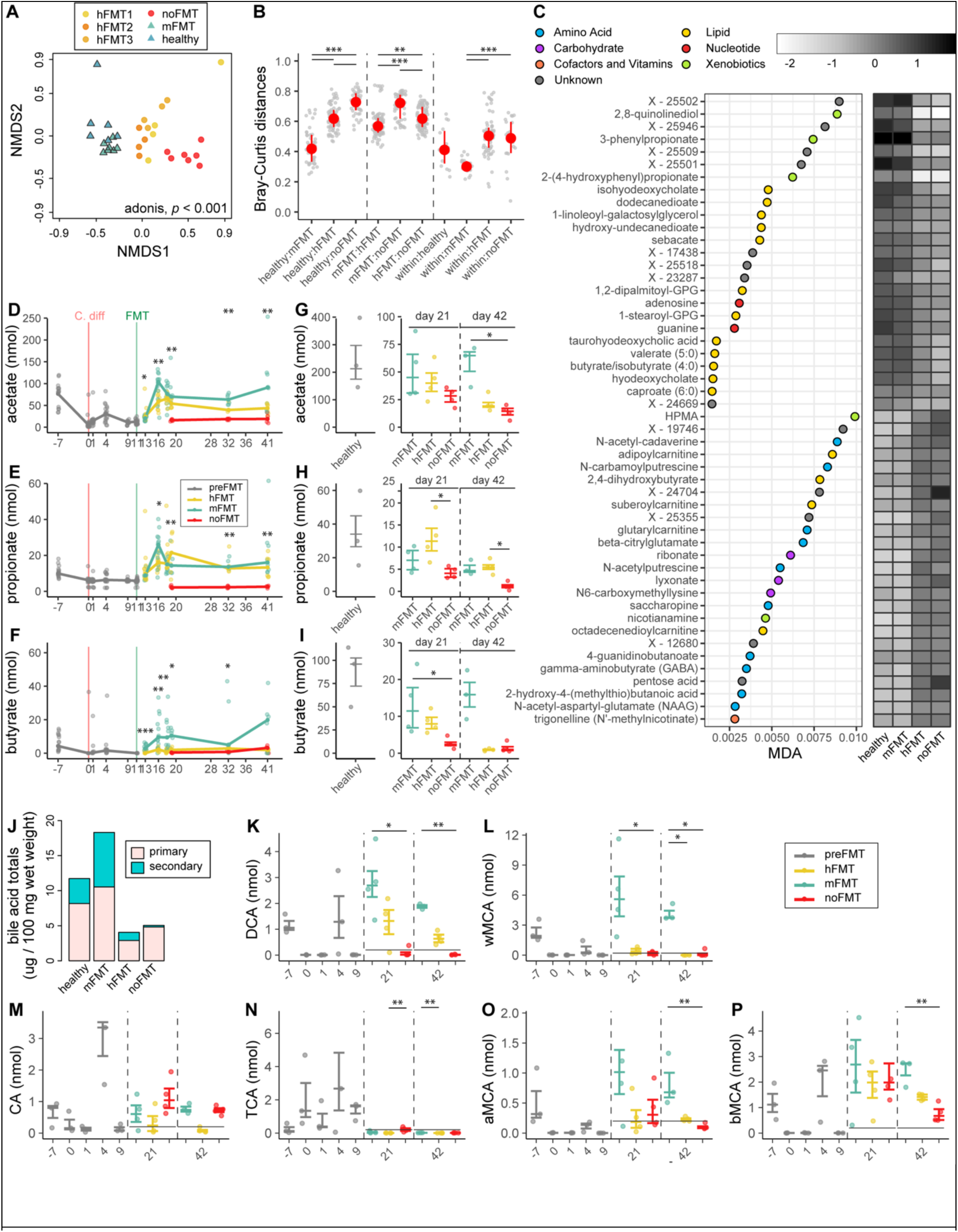
Mice treated with human feces (hFMT) demonstrate differential metabolic recovery. **A)** Non-metric multidimensional (NMDS) scaling of the Bray-Curtis dissimilarity index calculated from metabolite abundances from untargeted metabolomics in cecal samples of uninfected mice (healthy), mice with rCDI receiving no treatment (noFMT), or treated with human (hFMT1 – 3) or mouse feces (mFMT). PERMANOVA, *p* < 0.001 calculated across groups (healthy, mFMT, hFMT1 – 3 combined, and noFMT). **B)** Pairwise Bray-Curtis dissimilarity calculated between (healthy, mFMT, hFMT, and noFMT) and within groups. Statistical significance determined using Kruskal-Wallis test, with a post-hoc Dunn test (\**p* < 0.01, \*\**p* < 0.001, \*\*\**p* < 0.0001). **C)** Mean Decrease Accuracy (MDA) of top 25 important metabolites increased or decreased post-FMT from untargeted metabolomics, as identified by Random Forest Analysis. Mean log_10_-normalized median-scaled and minimum-imputed abundance of each metabolite. Targeted analysis of **(D)** acetate, **(E)** propionate and **(F)** butyrate fecal abundances over time in mice with rCDI treated with mFMT, hFMT, or noFMT. Targeted analysis of cecal **(G)** acetate, **(H)** propionate and **(I)** butyrate abundance at day 21 or 42 post-infection in uninfected mice (healthy; grey) compared to mice with rCDI treated with mFMT, hFMT, or noFMT. **J)** Total levels of primary and secondary bile acids from targeted metabolomics in the ceca of uninfected mice (healthy) compared to mice with rCDI treated with mFMT, hFMT, or noFMT.. **K)** Deoxycholic acid (DCA), **L)** μ-muricholic acid (MCA), **M)** cholic acid (CA), **N)** taurocholic acid (TCA), **O)** α-MCA and **P)** β -MCA in the same mice. Statistical significance for **(D – P)** determined using Kruskal-Wallis test, with a post-hoc Dunn test for **(G – I, K – P)** (\**p* < 0.05, \*\**p* < 0.005, \*\*\**p* < 0.0005).

We also conducted targeted analysis of metabolites previously associated with *C. difficile* susceptibility or resistance. The predominant gut SCFAs acetate, propionate, and butyrate all decreased during antibiotic treatment (Figure 5D-I). While acetate and propionate levels both increased following mFMT or hFMT compared to untreated mice, butyrate levels remained significantly decreased in mice given hFMT compared to mFMT (Kruskal-Wallis, *p* < 0.05). Of note, cecal levels of all three SCFAs never recovered to levels of uninfected mice, with untreated and hFMT-treated mice demonstrating the lowest levels of cecal butyrate (Figure 5G-I). Total primary and secondary bile acids were also decreased following hFMT compared to mFMT (Figure 5J). Following antibiotic exposure, we observed overall decreases in the secondary bile acids deoxycholic acid (DCA) and μ-muricholic acid (w-MCA), as well as the primary bile acids cholic acid (CA), α-MCA, and β -MCA (Figure 5K-P). Overall, levels of any measured bile acid were decreased in the hFMT-treated group. While partial recovery of DCA was observed in hFMT-treated mice (Figure 5K), little recovery of the mouse-specific w-MCA was observed in mice that did not clear (hFMT or noFMT) (Figure 5L). The primary bile acid and spore germinant, taurocholic acid (TCA), increased following antibiotic exposure, but decreased back to low levels in all three groups after treatment (Figure 5N). In contrast, cholic acid (CA) was highest in the no FMT group and lowest in the hFMT-treated group (Figure 5M).

## Discussion

The success of FMT in treating rCDI has been linked to both the restoration of key taxa (29, 30, 64) as well as the recovery of specific metabolic functions (24, 32, 34). Although specific OTUs after FMT vary across human studies, taxonomic similarity across individuals likely fulfills redundant and necessary functions that aid *C. difficile* clearance. Our results using a mouse model of rCDI suggest that engraftment alone does not guarantee the functional outcomes necessary for *C. difficile* clearance. Despite observing engraftment of similar taxa and recovery of many gene-encoded functions following both mFMT and hFMT mice, *C. difficile* clearance did not occur with hFMT treatment. The inability to clear *C. difficile* was accompanied by deficits in microbial metabolites typically associated with clearance, implying that the mere presence of ‘healthy’ taxa and their gene-encoded functions is not sufficient to ensure FMT success. While human studies suggest that inter-individual variation in FMT outcomes for rCDI may be of minimal concern, host-or microbe-related differences between recipients and their donors could explain FMT failures for rCDI or decreased efficacy for conditions other than rCDI (45, 69, 70).

Our study assessed microbiome recovery after FMT from human- or mouse-derived sources by examining microbial composition (taxonomic), genomic potential (gene-encoded functions), and metabolites (realized functions). At the compositional level, both 16S rRNA gene-based and metagenomic sequencing demonstrated increased recovery of overall diversity and species engraftment from hFMT into mice. These findings suggest that failure to clear *C. difficile* was not due to an inability of human microbes to colonize the mouse gut. This observation aligns with previous studies showing that a humanized microbiota can engraft antibiotic-treated mice (71) and specifically that human microbes in a mouse can confer resistance to primary infection with *C. difficile* (72, 73). However, *C. difficile* clearance in mice using a human microbiota in the context of rCDI has not been demonstrated in the literature. More commonly, studies have assessed the ability of different human feces to resist initial *C. difficile* colonization, using disease severity as a measure of *C. difficile* resistance to identify potentially relevant resistance mechanisms (74–76). This indicates that while human microbiota transplanted into mice may be effective in preventing the initial germination and colonization of *C. difficile*, clearing an established infection requires a different mechanism, one that, at least in our model, could not be achieved by human microbes transplanted into a mouse gut.

Compared to taxonomic differences, differences in the gene-encoded functions across the FMT groups in our study were less pronounced. In human studies of FMT for rCDI, metagenomic analyses have primarily focused on pre-versus post-FMT changes, comparing two markedly different gut environments. More nuanced comparisons that could identify microbial genes specific to CDI recovery, such as those between successful and failed cases, are challenging due to limited sample sizes (77, 78). Our metagenomic results demonstrate that hFMT resulted in restoration of relevant functional potential, with many differences attributed to the individual fecal source independent of *C. difficile* clearance. Notably, hFMT-treated mice exhibited similar or even elevated levels of gene-encoded functions previously associated with *C. difficile* clearance (29, 38, 79, 80), including high representation of amino acid modulation in both mFMT- and hFMT-treated mice compared to untreated mice. It is possible that this result is explained by inherent bias of commonly used databases, which include many unknown bacterial genes, many of which may be biased towards cultured organisms and human-associated bacteria (81, 82). For instance, the genes responsible for the transformation of muricholic acid, a mouse-specific secondary bile acid, remain unidentified. If this function is the mouse-specific representative of the *bai* operon, the operon responsible for the transformation of deoxycholic acid in humans (67), our metagenomic analyses would not recover this. While our findings highlight the successful transplantation of human-derived microbes in mice, they also underscore the limitations of our current metagenomic tools in fully capturing functional dynamics, particularly in mouse models. This highlights the need for more comprehensive databases that include a broader range of bacterial genes, especially from non-human backgrounds.

Our metabolomic analyses concurred with metabolomic analyses of patients with rCDI pre- and post-FMT, demonstrating recovery of SCFAs and bile acid ratios only alongside successful clearance (32, 34, 38). In our study, mice that received hFMT exhibited lower butyrate levels compared to mice that cleared, despite metagenomic presence. Mice treated with hFMT also displayed reduced levels of secondary bile acids compared to uninfected and mFMT-treated mice, despite the known genes being present in their metagenomic counterparts. Recent advances have identified new microbial-derived bile acids that may hold importance for CDI and other host-microbe interactions, which were not included in our targeted approach (83). Emerging research has also highlighted factors beyond BA modulation and SCFA production that may aid CDI recovery, such as nutrient competition in a resistant microbiota (84, 85). *C. difficile* has demonstrated metabolic flexibility and is capable of metabolizing various amino acids (40). Colonization by a microbiota with diverse amino acid-utilizing capabilities is thought to restrict *C. difficile* through depleting available free amino acids (36, 37, 86). Although we did not directly measure amino acid levels, we observed distinct taxa across FMT groups that contribute these functions.

While transplanting microbes from one host to another represents an extreme comparison, it is reasonable to posit that certain host environments (inclusive of their extant microbes and environmental or host-specific differences) may not be translatable across the human population or disease conditions. A recent study suggested that the success of FMT in treating rCDI depends on donor-derived species that initially reduce inflammation through metabolite production, thereby facilitating the recovery of existing recipient microbes (87). Variations in diet, which has been demonstrated to influence severity of CDI (88, 89), could also drive differences in how a transplanted microbial community behaves in another host, whereby adaptation to the host’s previous diet no longer renders the same function in the new host. Perhaps most relevant, the immune status of the recipient may drive differential recognition, and thus variable activity, of the transplanted microbiota (90). Additionally, interactions between host and microbiota, including immune response, likely impact FMT efficacy. Prior inflammation has been demonstrated to increase CDI severity in mice via sustained presence of pro-inflammatory Th17 cells (91). More relevant to clearance, it was observed that *Rag1^-/-^* mice had a decreased capacity to clear *C. difficile* in a model of primary CDI (41). While our study did not assess immune profiles, species-specific host differences to recognize their microbial ‘self’ might impair the collective functional ability of the microbiome (92). Finally, interactions with other species may influence *C. difficile* virulence and microbiota functions, as has been demonstrated with *Enterococcus* (43). Altogether, this highlights the complexity of ecological factors involved in microbe-mediated conditions, many of which remain unresolved. Our findings provide additional insight into host-microbe interactions, offering potential avenues for enhancing the effectiveness of microbial interventions.

## Acknowledgments

Thank you to our benevolent participants for their fecal donations. We would like to acknowledge Clemson University for generous allotment of compute time on Palmetto cluster. We also thank Kwi Kim at the University of Michigan for helping us with the fecal HPLC data. This publication was made possible, in part, with support from the Clemson University Genomics and Bioinformatics Facility, which receives support from an Institutional Development Award (IDeA) from the National Institute of General Medical Sciences of the National Institutes of Health under grant number P20GM109094. AMS was supported by grant number K01-DK111794 from the National Institute of Diabetes and Digestive and Kidney Diseases. VBY was supported by AI124255 and AI090871.

S.M. – Data Curation, Formal Analysis, Investigation, Methodology, Software, Writing – original draft; and Writing – review and editing; K.C.V. – Data Collection and Curation, Methodology– original draft and Writing – review and editing; V.B.Y. – Supervision, Project Administration, Funding Acquisition – original draft, and Writing—review and editing; A.M.S. – Conceptualization, Data Collection and Curation, Formal Analysis, Methodology, Investigation, Supervision, Project Administration, Funding Acquisition, Writing – original draft and Writing – review and editing.

## Supplemental figure legends

**Figure S1.**
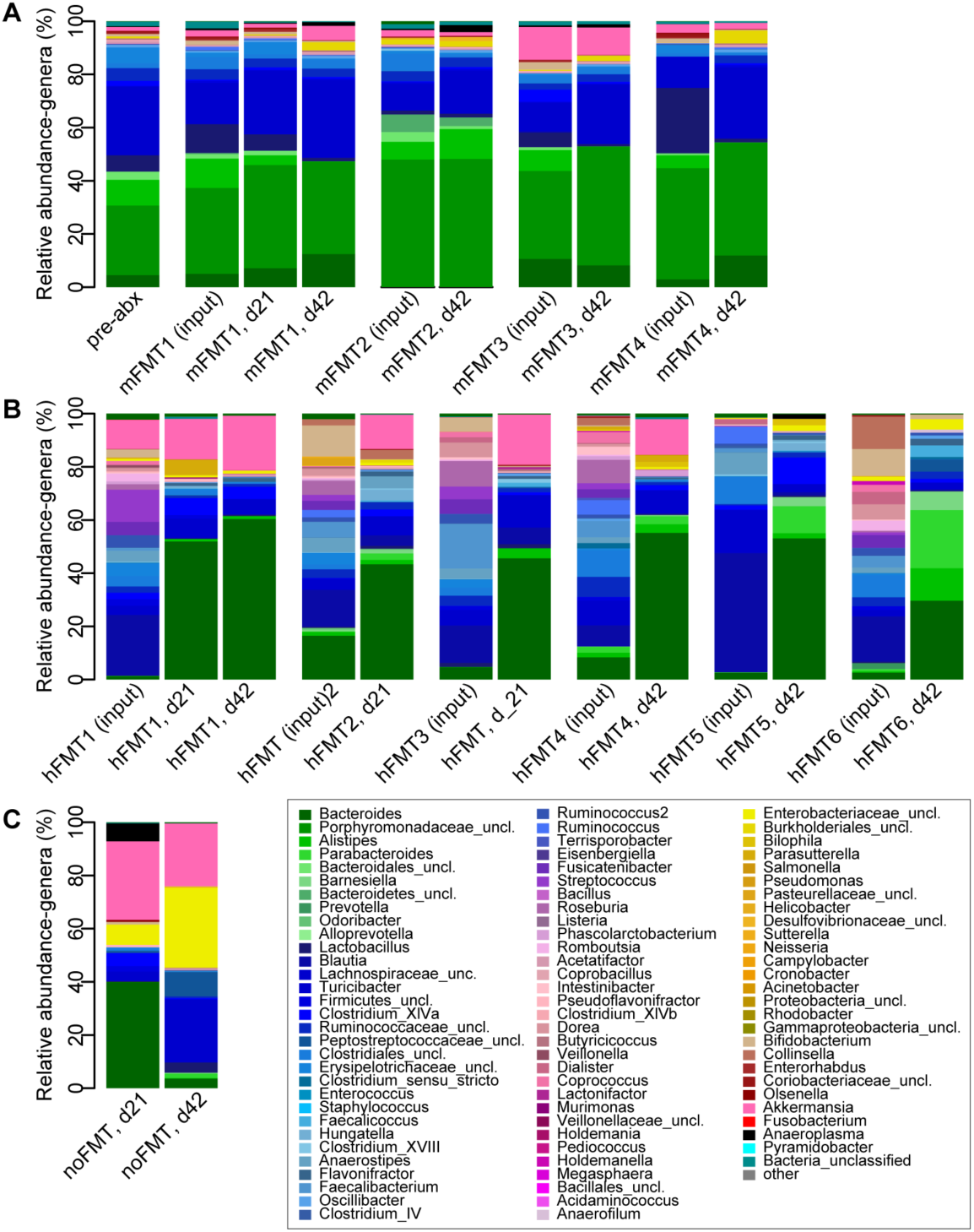
Genus-level taxonomy across mice by donor type. Average relative abundance of top 98% genera observed in FMT inputs and cecal samples in mice **A)** prior to any treatment (pre-abx) or after different individual mFMT sources, **B)** different individual hFMT sources, or **C)** no FMT, at indicated timepoints (day 21 or 42 post-infection).

**Figure S2.**
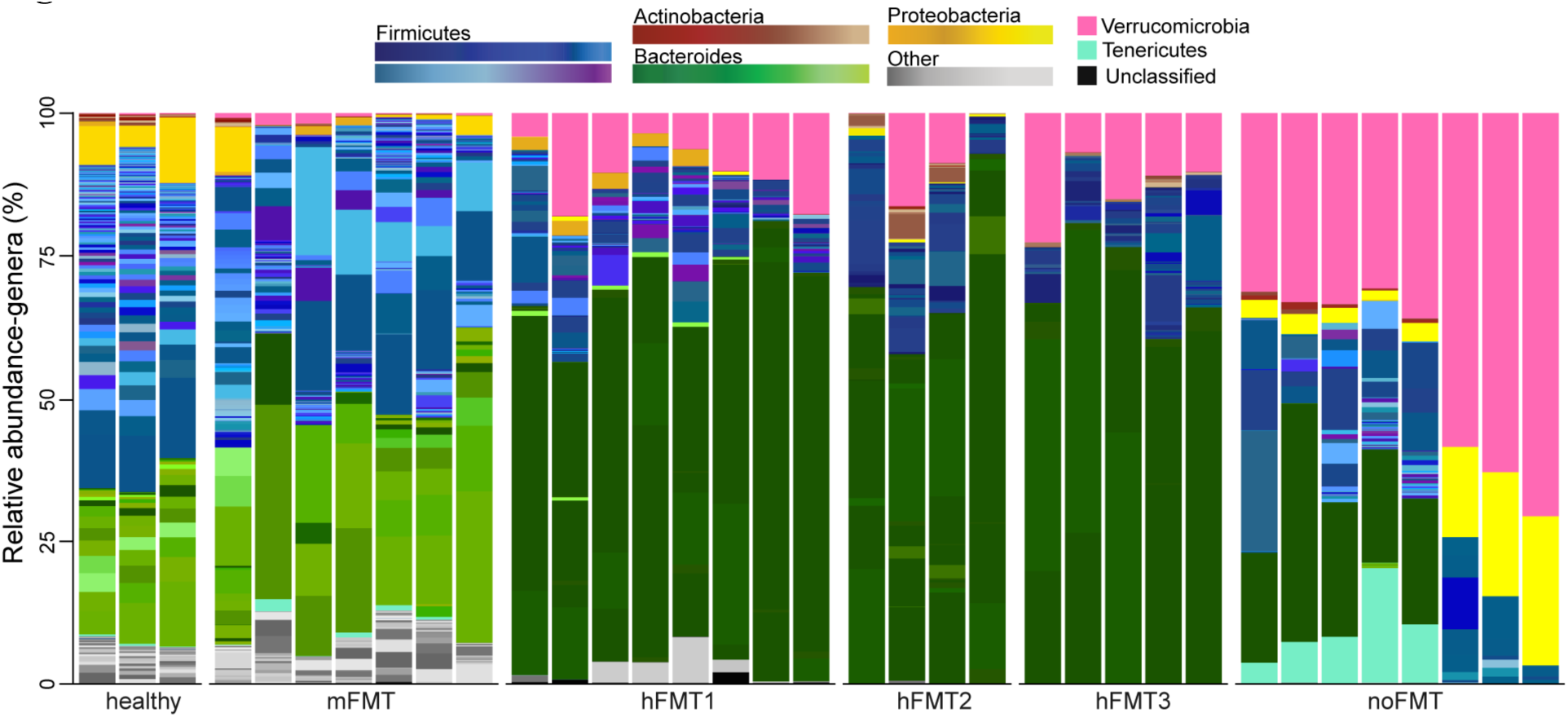
Relative abundance of individual mice. Relative abundance of genera identified by MetaPhlAn4 from cecal samples of mice. Each bar represents an individual mouse.

**Figure S3.**
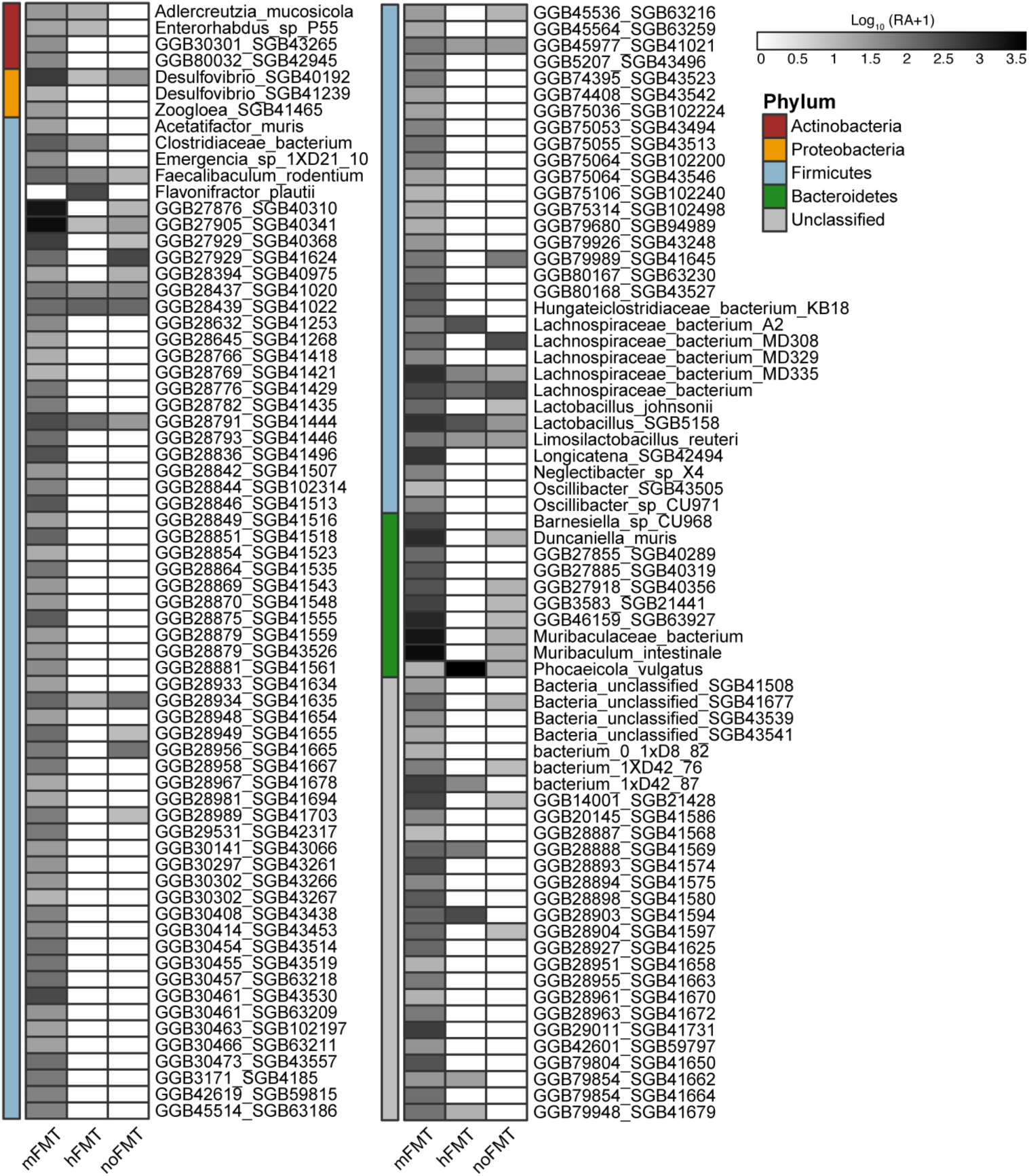
Bacterial genera significantly different in abundance between cleared and colonized mice. Mean log_10_-transformed (CPM + 1) of all bacterial genera significantly different in abundance between mice that cleared (mFMT) or did not clear (hFMT, noFMT) *C. difficile* (based on MaAsLin2; linear model with BH correction, *q* ≤ 0.01).

**Figure S4.**
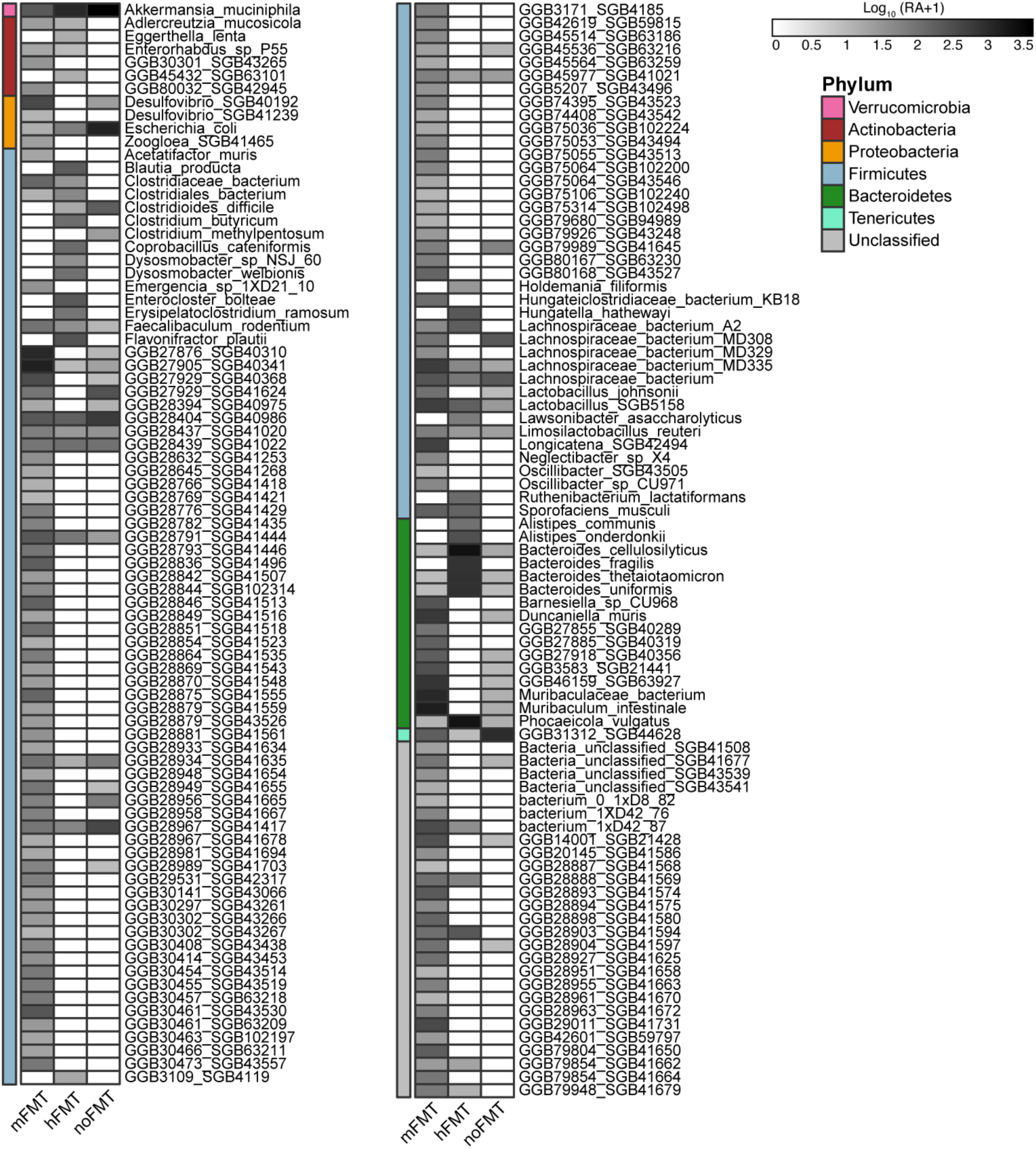
Bacterial genera significantly different in abundance across treatment groups. Mean log_10_-transformed (CPM + 1) of all bacterial genera significantly different in abundance across treatment groups compared to mice treated with mFMT (based on MaAsLin2; linear model with BH correction, *q* ≤ 0.01).

**Figure S5.**
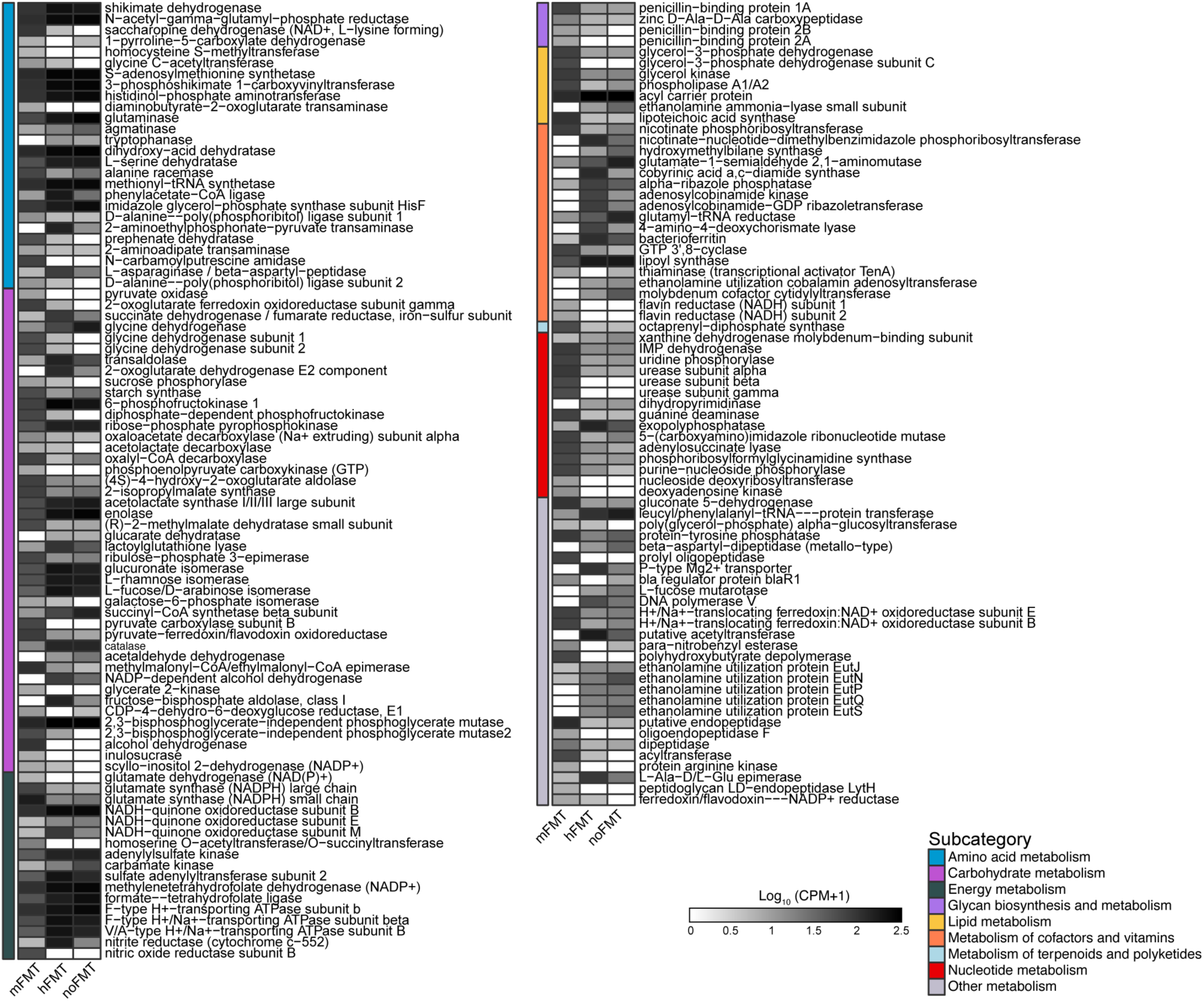
Microbial genes (KOs) associated with metabolism significantly different in abundance between cleared and colonized mice. Mean log_10_-transformed (CPM + 1) of metabolism associated KEGG orthologs (KOs) significantly different in abundance between mice that cleared (mFMT) or did not clear (hFMT, noFMT) *C. difficile* (based on MaAsLin2; linear model with BH correction, *q* ≤ 0.001).

**Figure S6.**
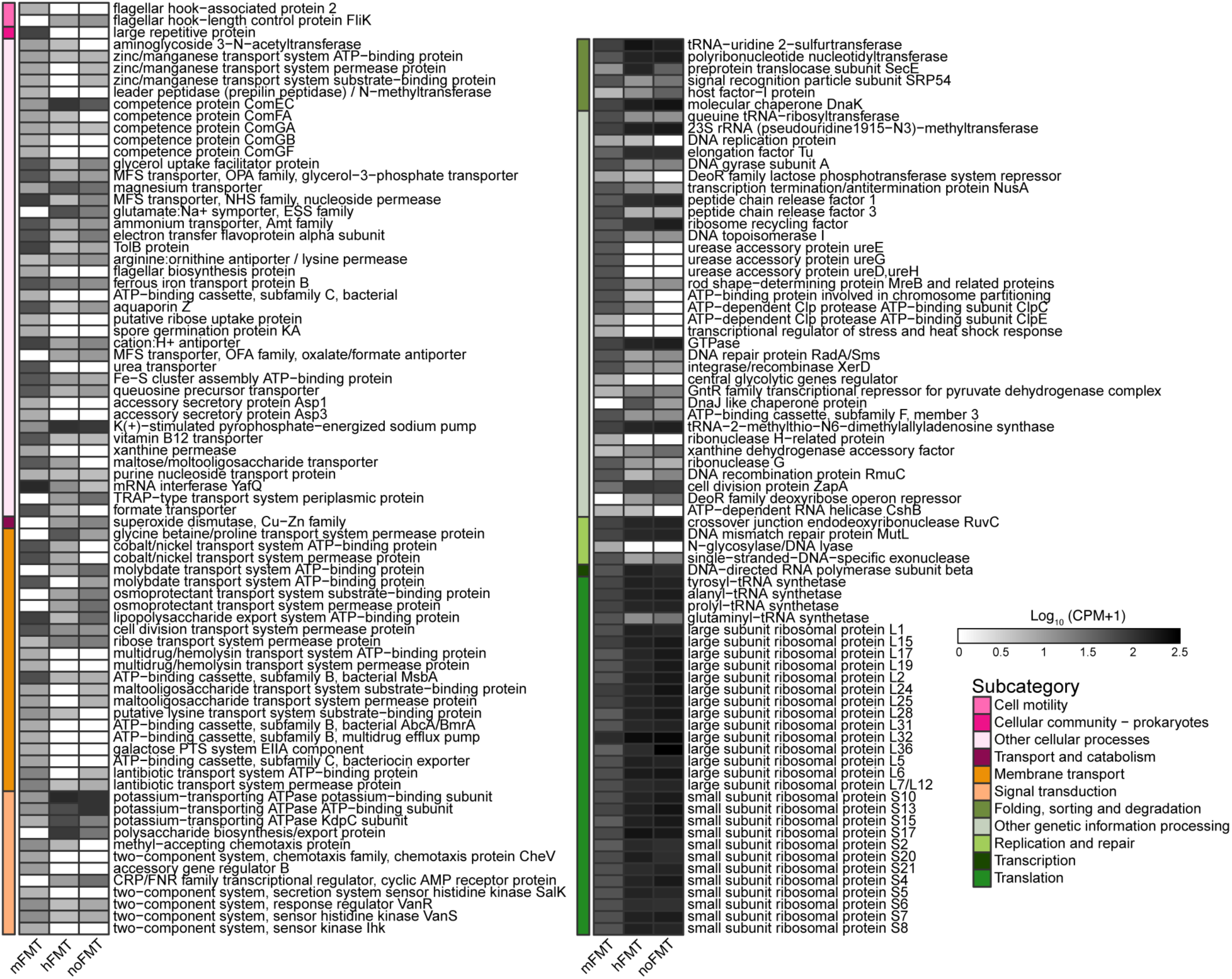
Microbial genes (KOs) not associated with metabolism significantly different in abundance between cleared and colonized mice. Mean log_10_-transformed (CPM + 1) of Kos from non-metabolism categories significantly different in abundance between mice that cleared (mFMT) or did not clear (hFMT, noFMT) *C. difficile* (based on MaAsLin2; linear model with BH correction, *q* ≤ 0.001).

**Figure S7.**
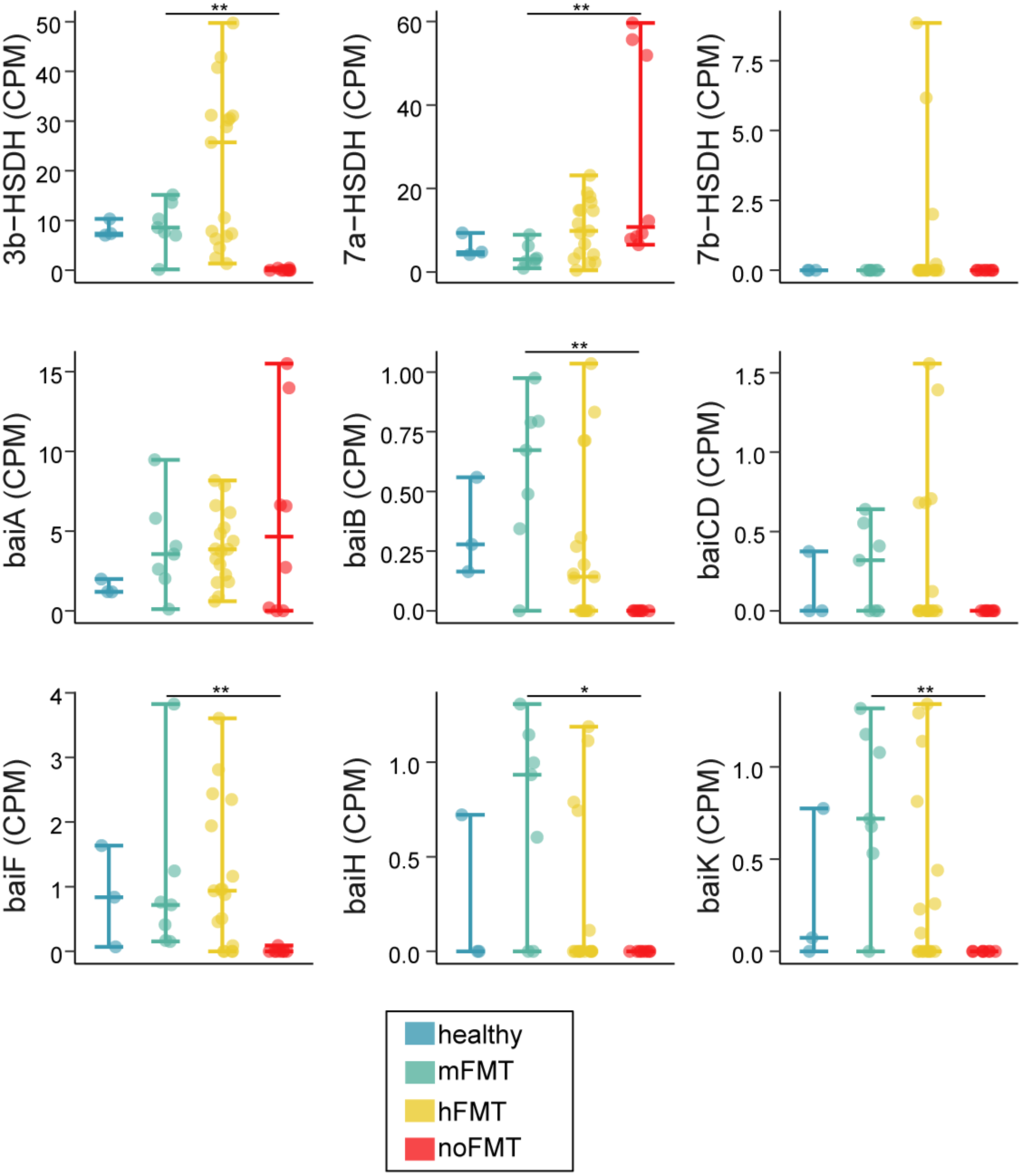
Abundance of microbial genes associated with bai operon. CPM of Uniref90 genes associated with the bile acid inducible (bai) operon. Statistical significance determined using Kruskal-Wallis test, with a post-hoc Dunn test (\**p* < 0.01, \*\**p* < 0.001)

**Figure S8.**
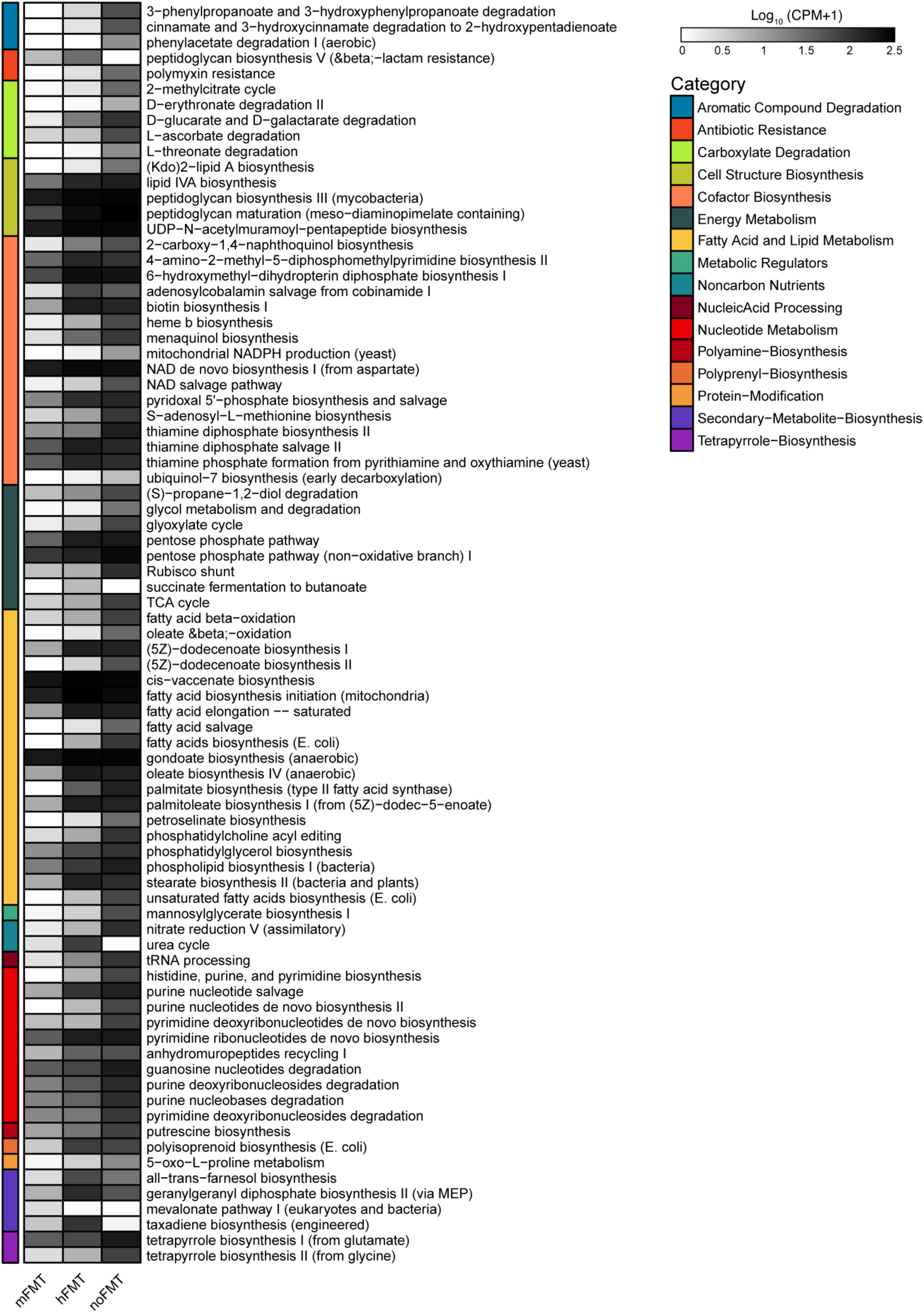
Microbial pathways associated with other metabolism significantly different in abundance across treatment groups. Mean log10-transformed (CPM + 1) of other metabolism MetaCyc Pathways significantly different across treatment groups compared to mice treated with mFMT (based on MaAsLin2; linear model with BH correction, *q* ≤ 0.001).

